# Genetic depletion of BDNF impairs both context-dependent and context-independent extinction learning of a spatial appetitive task

**DOI:** 10.1101/2021.01.26.428221

**Authors:** Marta Méndez-Couz, Denise Manahan-Vaughan

## Abstract

Brain derived neurotropic factor (BDNF) supports neuronal survival, growth, and differentiation and is involved in forms of hippocampus-dependent learning, as well as hippocampus-dependent learning. Extinction learning (EL) comprises active inhibition of no-longer relevant learned information, in conjunction with a decreased response of a previously learned behavior. It is highly dependent on context, and evidence exists that it requires hippocampal activation. Concordantly, the participation of BDNF in hippocampus-dependent memory is experience-dependent. BDNF has been associated with synaptic plasticity needed for acquisition and extinction learning of fear conditioning. However, little is known about its influence on the extinction and renewal of spatial appetitive extinction learning (EL). In this study, in BDNF^+/−^-mice we evaluated to what extent BDNF contributes to spatial appetitive EL in the presence (ABA) or absence (AAA) of a context change. Daily training, to reach acquisition criterion in a T-maze, resulted in a similar outcome in BDNF^+/−^-mice or their wildtype (wt) littermates. EL was delayed in the AAA, and significantly impaired in the ABA-context compared to EL in wt littermates. When renewal was tested in the ABA paradigm we detected a significant response in wt controls, but not in BDNF^+/−^-mice. Taken together, these results support an important role for BDNF in EL in AAA and ABA context, as well as renewal of a spatial appetitive task, processes that relate to information updating and retrieval.

## 1. INTRODUCTION

Operant behaviors are voluntary actions controlled by their consequences. Animals readily acquire behaviors (e.g., lever pressing) to obtain a desirable outcome (e.g., food pellet or drug delivery) and equally learn to suppress or diminish this behavior when the reinforcer is withdrawn, in a process known as extinction learning (EL) (Eddy et al., 2016). EL of instrumental responding is a central point of behavioral change (Todd et al., 2014; Bouton, 2019). In opposition, behavior, the associated responses for which have been extinguished, can re-emerge through several mechanisms. One of these processes is referred to as renewal, which occurs in the form of a return of the associated response, when an animal is tested in a context different from the one in which extinction learning took place (Bouton and Bolles, 1979). The latter is referred to as an ABA paradigm. This reappearance of responding demonstrates that EL does not comprise erasure of the original learning. In this context, EL may be considered to be a new form of learning, involving memory formation whilst preserving the original memory trace.

Although, in mechanistic studies of EL the primary focus has been placed on aversive forms, studies on appetitive forms of EL in rodents have offered novel insights as to the cognitive challenges, brain structures and molecular systems involved in this process. Where EL of a spatial appetitive task was studied, it was found that EL in the absence of a context change requires many more task exposures and increased catecholaminergic contribution compared to context-dependent EL (Andre et al., 2015b). EL in the absence of a context change also depends on activation of the metabotropic glutamate receptor, mGlu5, whereas context-dependent EL does not (Andre et al., 2015a). Context-dependent EL recruits information processing in the hippocampus that involves gene encoding (Mendez-Couz et al., 2019), a process that has been linked to hippocampal synaptic plasticity (Kemp et al., 2013) and to context-dependent spatial learning (Hoang et al., 2018).

One very important mediator of signaling pathways related to gene encoding and synaptic plasticity in the brain is brain-derived neurotrophic factor (BDNF). It supports neuronal survival as well as functional and structural synaptic plasticity (Zagrebelsky and Korte, 2014). It has been proposed that activity-dependent secretion of BDNF may support synapse-specific synthesis of proteins that are required for the stability of long-term forms of synaptic plasticity (Lucidi-Phillipi et al., 1995; Poo, 2001; Novkovic et al., 2015). Moreover, secreted BDNF is capable of mediating many activity-dependent processes in the mammalian brain, including neuronal differentiation and growth, synapse formation and plasticity, and higher cognitive functions (Park and Poo, 2013), including spatial learning (Aarse et al., 2016). This is especially notable, given that synaptic plasticity in the hippocampus, in the forms of long-term potentiation and long-term depression, comprises the primary cellular mechanism underlying long-term spatial memory (Kemp and Manahan-Vaughan, 2007).

BDNF has been strongly implicated in clinical depression and cognitive impairments observed in depressed patients (Brunoni et al., 2008). In fact, BDNF levels can be significantly influenced by exposure to stress (Bath et al., 2013), a process that may take place during fear conditioning learning and aversive forms of extinction learning. In line with this, BDNF contributes to EL of fear conditioned memory (Peters et al., 2010; Rosas-Vidal et al., 2018; Notaras and van den Buuse, 2020). Specifically, the role of BDNF has been scrutinized in studies of malfunctioning contextual fear conditioning and disrupted EL, where BDNF was infused into the hippocampus (Kirtley and Thomas, 2010) or the infralimbic cortex (Peters et al., 2010). Antagonism of BDNF in the prefrontal cortex in an animal model of aversive EL results in impairment of EL recall and changes in ventral levels of BDNF (Rosas-Vidal et al., 2018). In humans, the genetic BDNF^Val66Met^ variant is associated with impaired EL (Notaras and van den Buuse, 2019, 2020) as well a decreased response for fear-extinction therapies (Soliman et al., 2010).

Although the role of BDNF in EL of conditioned aversive learning has been well-described (Karpova et al., 2011; Psotta et al., 2013), little is known about the role of BDNF in *appetitive* EL of context-related experience. Here, we studied EL of a spatial appetitive task in the presence or absence of a context-change, in BDNF^+/−^ mice. We report no deficiencies in task acquisition in the transgenic mice compared to their wildtype littermates. However, EL in the absence of a context change (AAA paradigm) was delayed in BDNF^+/−^ mice compared to their wt littermates. Furthermore, context-dependent EL was significantly impaired and renewal was absent in the ABA paradigm. These findings support a role for BDNF in information updating related to EL and renewal under spatial appetitive circumstances.

## 2. MATERIAL AND METHODS

This study was carried out in accordance with the European Communities Council Directive of September 22, 2010 (2010/63/EU) for care of laboratory animals. All experiments were performed according to the guidelines of the German Animal Protection Law and were approved by the North Rhine-Westphalia State Authority (Landesamt für Arbeitschutz, Naturschutz Umweltschutz und Verbraucherschutz, LANUV, Bezirksamt, Arnsberg). Animal numbers were kept to the minimum required for biometric planning.

BDNF^+/−^ mice and their wild-type (WT) littermates were used (Animal breeding facility of the Medical Faculty, Ruhr University Bochum). The strain was originally established by Korte and colleagues (Korte et al., 1995), whereby one allele of the BDNF protein-coding exon was replaced by a neomycin-resistant gene surrounded by a glycerate kinase gene promotor and a polyadenylation signal. Most of the mature coding sequence of BDNF was deleted through this replacement and the inserted neomycin resistant gene serves as a biomarker. Heterozygous BDNF^+/−^ mice, and their wt littermates, were produced by crossing BDNF^+/−^male mice with C57BL/6 female mice. Homozygote BDNF^-/-^ mice were not used as they express abnormalities including neuronal loss and retarded development (Korte et al., 1995; Zagrebelsky and Korte, 2014).

### Behavioral paradigm

The experimental room was faintly illuminated during experiments and animal behavior was recorded by means of a monitoring system (Videomot; TSE Systems, Bad Homburg, Germany) that enabled subsequent offline analysis. EL experiments were conducted in a T-maze that was composed of a starting box (20cm **×** 20 cm) that was separated from the main corridor (100 cm **×** 20 cm) by a sliding door and two side corridors (40 cm **×** 10 cm) positioned perpendicularly to the other end of the main corridor (Figure 1). The walls were 40 cm high. In each side corridor, a small round cup was placed 1 cm in front of the end wall and in the middle of the floor, where the reward was hidden from distant view. The reward consisted of 100 μl of 25% sucrose solution. This form of reward was chosen because according to Adachi and colleagues, the deletion of BDNF in the hippocampal CA1, or DG region does not alter preference for a natural reward in the sucrose preference test in comparison to controls (Adachi et al., 2008). The maze design and the protocol followed were as described previously (Andre et al., 2014; Wiescholleck et al., 2014; Andre and Manahan-Vaughan, 2015) (Figure 1).

**Figure 1:**
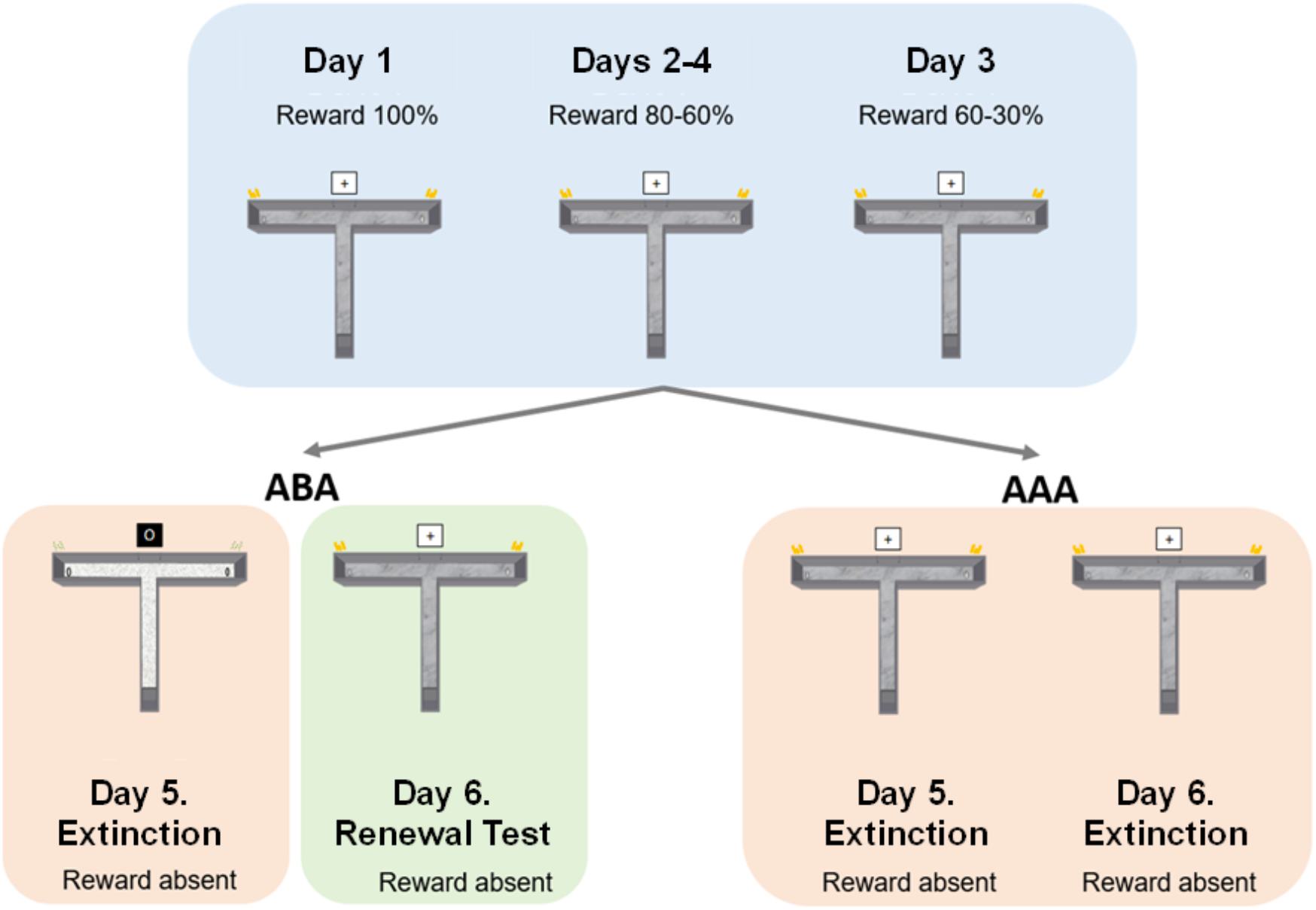
Schema of the behavioral protocol. Top: On Day 1, animals participate in two sessions each comprising ten consecutive trials each (separated by 10-min intervals) that include a reward probability of 100%. On Day 2 two ten trial sessions at 80% reward probability occur. On Day 3, the reward probability of the first session is 80%, and of the second session is 60%. On Day four, the reward probability declines from 60% in the first session, to 30% in the second session. **Bottom right**: On Days 5 and 6, animals participate in the extinction learning protocol in the same context (AAA) in the absence of a reward. **Bottom left**: In the ABA paradigm animals engage in EL in the presence of a context change on Day 5. During extinction learning, the context (floor pattern, distal cues, odor cues) are changed. On day 6, renewal is tested by re-exposure to context A. No reward is present in the maze during the extinction or renewal test days.

The context of the maze was changeable by three aspects, floor pattern, external visual cues, and internal odor cues. The maze floors had distinct visual patterns such as wood-like, totally white, or granite-like patterns printed on washable PVC. During each phase of training or testing, 1 µl of a particular odor (e.g. almond or vanilla food aroma, Dr. Oetker, Bielefeld, Germany) was placed at the end of the two arms. The odor remained consistent throughout all trials, with the exception of context-dependent EL, where the odor was changed. For visual distal cues, Din A4 size cards of white paper printed with a thick black cross, or a black-filled circle, were used. These visual cues were positioned 40 cm above the end of the main corridor (Figure 1). The visual cues remained constant in all trial conditions with the exception of EL in the ABA paradigm.

To test context-independent EL in the T-maze an AAA paradigm was used, in which training and all of the extinction sessions were conducted in the same context (Figure 1). Here, EL was followed on two consecutive days. To examine context-dependent EL we used an ABA paradigm, in which training was conducted in Context A, while the EL session was conducted in Context B (Figure 1). Renewal was subsequently tested in context A.

A week before the beginning of the behavioral training, the mice were weighed and food availability was reduced to achieve a minimal weight of 85% of the previously determined body weight. This weight was sustained until the end of the experiment.

Every day, each mouse underwent a learning session consisting of 2 sessions of 10 consecutive trials with an intertrial time of 15 seconds and half an hour difference in between sessions. Each trial began when the door to the starting box was opened and the animal could enter the maze. It ended when the animal entered an arm of the T-maze or when 30 s had elapsed without leaving the starting box. In each trial, the animals were expected to search for a reward that was placed in the indentation located in the floor at the end of a predetermined arm. From Days 1 through 4, the reward probability was decreased stepwise from 100 to 30% to increase extinction resistance. Without this form of training, testing contextual changes during repeated extinction trials in the absence of a food reward would not have been possible (Andre et al., 2015a). Learning criterion was reached when the animal successfully entered the correct arm on eighty-five per cent of the trials in the last session. Animals that did not reach a minimum of 85% correct responses by the final trial of Day 4, were excluded from the study. In between trials, the maze was wiped with a humid cloth to mix the odor trail that the animal could have left behind. Before the next animal was exposed to the T-Maze it was thoroughly cleaned and dried.

In the case of the ABA paradigm, extinction learning was evaluated on Day 5. For this purpose, mice were introduced into the T-maze in absence of food reward for 2 sessions of 10 trials during which the context (floor, odor, and cue card) was changed. On Day 6, renewal (RN) was assessed. Here, the animal was reintroduced to the original T-maze context (A) for another two sessions of ten trials each, but a food reward was also absent in this case.

In case of the AAA paradigm, mice underwent the same procedure in the original context (A), but during the 5^th^ and 6^th^ days, when EL was examined, the food reward was perpetually absent.

### Statistical Analysis

The resulting video recordings from each experimental phase were analyzed for the correct or incorrect response of the animals in each trial. To prevent bias, animals were coded so that the experimenter was unaware of the behavioral condition for each video analyzed. A two-way repeated measures (rm) analysis of variance (ANOVA) was used to determine the difference in between groups in the percentage of correct response for the acquisition and the extinction-renewal phases, with ‘group’ and ‘session’ as factors. Post-hoc analysis was conducted using a Holm-Sidak test, to determine the differences between individual trials. A one-way rm ANOVA were applied within-groups to test the effectiveness of EL and renewal, with ‘session’ as factor. Here, the Student-Newman-Keuls test was applied as post-hoc test. Sigmastat 11(Systat) and Prisma 8 (GraphPad) software were used for the analysis.

## 3. RESULTS

### A lack of BDNF does not impair context-related appetitive learning

No learning differences were found along the acquisition phase between BDNF^+/−^mice (n= 16) and their WT littermates (n= 16) (F_(1,210)_ = 0.25, *p* = 0.62). Animals exhibited equivalent improvements in choice behavior between sessions (F_(7,210)_ = 22.84, *p* < 0.01) (Figure 2). By day 4, both animal cohorts had reached the learning criterion of 85% correct choices.

**Figure 2:**
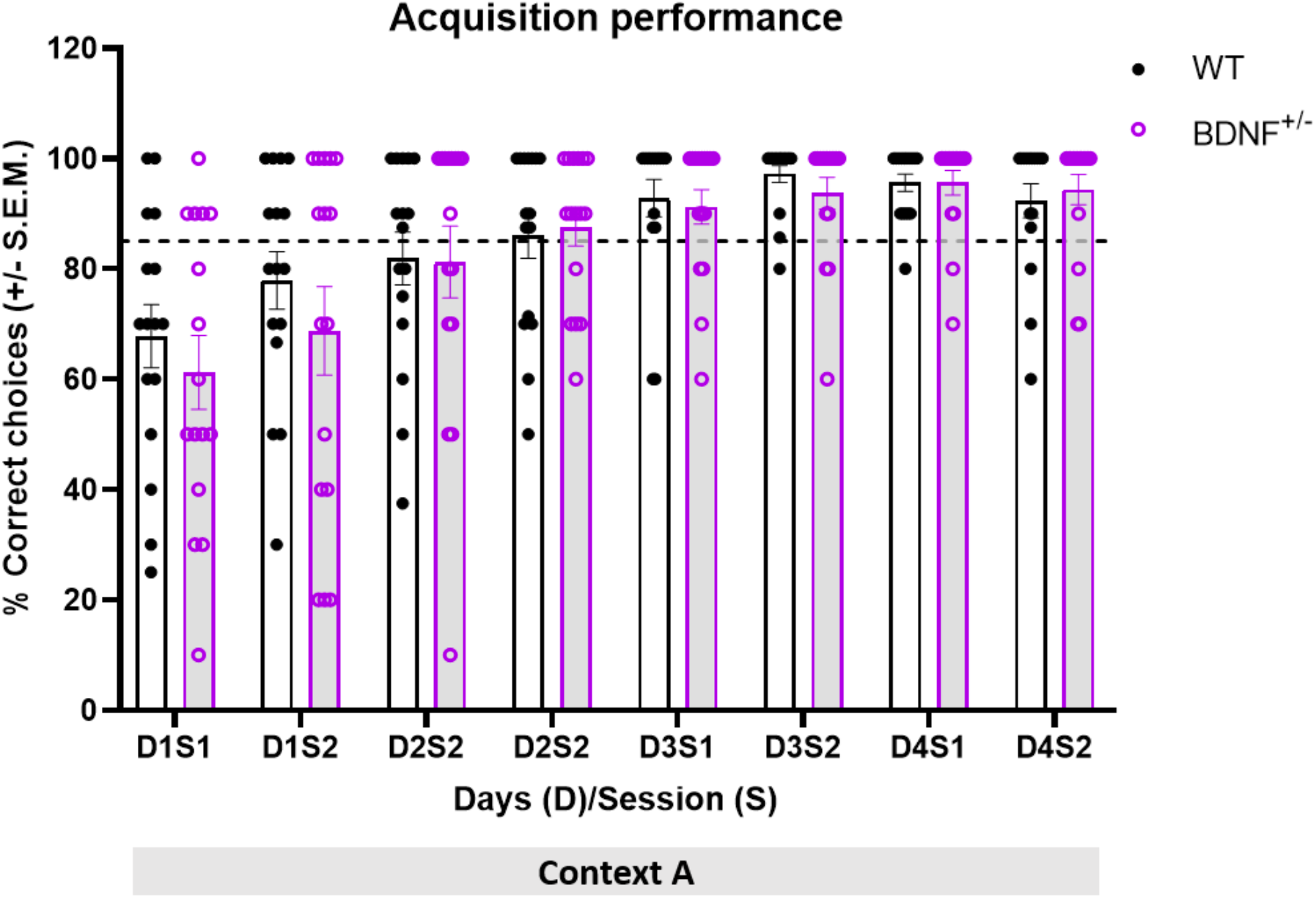
BDNF^+/−^ mice do not exhibit impairments in context-dependent task acquisition. The bar charts show the percentage of correct choices made by the animals during task acquisition learning over three days in context A. Each bar pair shows the animals’ correct choice performance in a given session (S) on a specific day (D) of acquisition. Black-outlined bars show the response of wildtype (WT) mice and purple-outlined bars show the response of transgenic BDNF^+/−^ mice. The dots (black: WT, purple: BDNF^+/−^) show the distribution of choice performance in a specific session. Both WT (n=16) and BDNF^+/−^ animals (n=16) acquired the task over the 3 day period, as signified by the increasing percentage of correct responses in both groups. Despite the gradual reduction in the reward probability, the two groups reached the 85% criterion of correct responses (dashed line) by day 4.

### A lack of BDNF is associated with a delay in extinction learning in the absence of a context change

To study the occurrence of extinction learning, we compared the last session of acquisition to the EL sessions on day 4 and day 5 in BDNF^+/−^mice (n= 8) with the performance of their WT littermates (n= 8). Both groups engaged in EL, as demonstrated by the differences between the final acquisition session (D4S2) and the EL sessions (F_(1,54)_ = 8.07, *p* < 0.01). Effects became apparent on the first day of EL in WT mice (Figure 3A) (wt D4S2 vs D5S2: F_(4,28)_ = 8.33, *p* < 0.01; Student Newman-Keuls *p* = 0.02), whereas the BDNF^+/−^ group did not show EL on the first EL day. By the second day of EL sessions, BDNF^+/−^mice exhibited EL that was significant compared to the last acquisition day session (BDNF^+/−^D4S2 vs D6S1: F_(4,28_) = 11.09, *p* < 0.01; Student Neuman Keuls *p* = 0.01) and EL performance was equivalent to levels achieved in wt mice.

**Figure 3:**
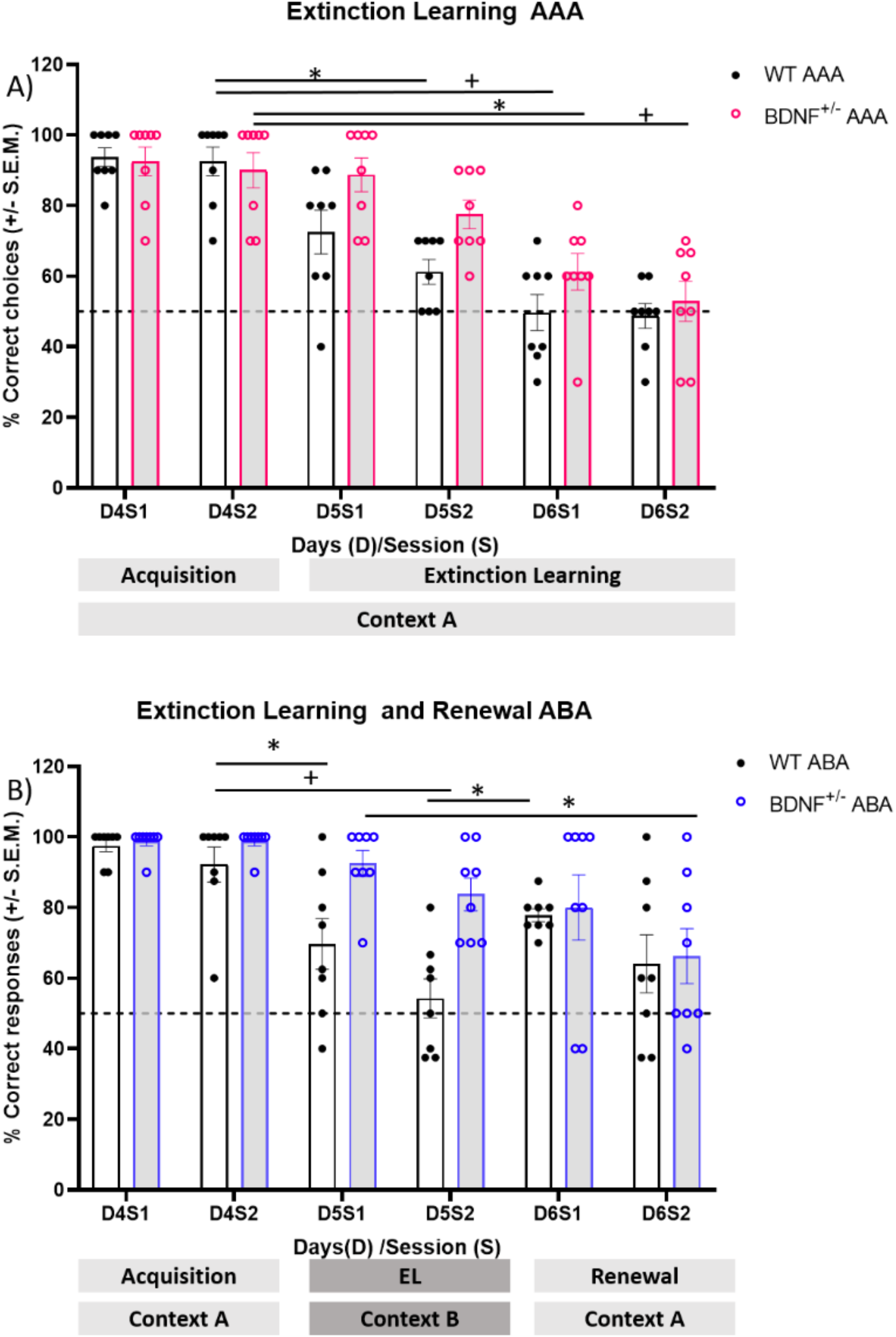
BDNF+/-mice exhibit a delay in extinction learning in the AAA context and an impairment in extinction learning in the ABA context. Renewal is impaired. The bar charts show the percentage of correct choices made by the animals during task acquisition on Day 4 and sessions 1 and 2 (D4S1, D4S2) in context A and during extinction learning over 2 days involving two training sessions per day (D5S1-D6S2) in context A. Each bar pair shows the animals’ correct choice performance. Black-outlined bars show the response of wildtype (WT) mice and color-outlined bars show the response of transgenic BDNF^+/−^ mice. The dots (black: WT, color: BDNF^+/−^) show the distribution of choice performance in a specific session, and the dashed line represents the chance performance level. **A) AAA paradigm**: Although correct choices in BDNF^+/−^ mice (n=8) and their WT littermates (n=8) were equivalent during both acquisition learning sessions on Day 4 (D4S1, D4S2), when extinction learning (EL) was assessed on Day 5, a significant difference between EL performance in WT and BDNF^+/−^ mice became apparent. WT mice showed rapid EL (D5S2, D6S1), whereas effects in BDNF^+/−^ mice were significantly slower. By end of Day 6, BDNF^+/−^ mice exhibited EL that was not significantly different from WT littermates, performing at chance levels (dashed line). B) In the ABA paradigm, BDNF^+/−^ mice (n=8) and their WT littermates (n=8) also displayed equivalent levels of correct choices during both acquisition learning sessions in context A on Day 4 (D4S1, D4S2). However, when EL was tested in context B on Day 5, a significant impairment in EL became evident in BDNF^+/−^mice compared to WT littermates (whereas WT mice exhibit performance differences between the end of acquisition, D4S2 to D5S1 and D5S2, BDNF^+/−^ do not). Although WT mice exhibited significant renewal on Day 6 (correct choices during D6S1 compared to choices during D5S2), BDNF^+/−^ mice displayed choice levels during D6S1, that were not different from choices made on Day 5, indicating that WT exhibited renewal and BDNF^+/−^ mice did not. During the second session on Day 6 (D6S2), WT animals displayed a decline in choosing the now unrewarded target arm, consistent with EL of the renewal response having occurred. BDNF^+/−^ mice displayed responses during this session (D6S2) that were equivalent to responses in D6S1 and D5S2. *(p≤0.05) ^+^(p≤0.01)

### A lack of BDNF is associated with impaired context-dependent extinction learning and an absence of renewal effects

Within the ABA paradigm, EL was significantly impaired in BDNF^+/−^mice compared to WT littermates (Figure 3B) (F_(1,56)_ = 9.64, *p* = 0.003). WT animals exhibited rapid and potent EL compared to the final acquisition trials of Day 4 (Figure 3B) (ANOVA F_(7,28)_ = 5.01, *p* = 0.004). In addition, significant differences occurred between the last session of acquisition and the first EL session in WT mice (wt D4S1 vs D5S1, Holm-Sidak post-hoc tes*t, p* = 0.019) and between the end of acquisition and the final EL session (wt D4S1 vs D5S2, Holm-Sidak post-hoc tes*t, p* < 0.001).

By contrast, BDNF^+/−^mice exhibited no EL in the first EL session (D5S1) compared to the last acquisition session (D4S2) (Figure 3B) (Holm-Sidak post-hoc tes*t, p* = 0.402). By the second set of EL trials on Day 5 (D5S2) effects were marginally non-significant compared to the final set of acquisition trials (D4S2) (Holm-Sidak post-hoc tes*t, p* = 0.051) (Figure 3B). Furthermore, EL performance was significantly different from wt mice (ANOVA F_(7,28)_=5.75, *p* = 0.002).

When renewal effects were assessed on Day 6 in the ABA cohort (Figure 3B), WT mice exhibited significant renewal during the first session compared to the final EL trial on Day 5)(D6S1 vs D5S2, Holm-Sidak, *p* = 0.015). Effects declined by the second renewal session on Day 2, consistent with an EL effect due to the absence of food reward (Figure 3B)(D6S2 vs D5S2, Holm-Sidak, *p* = 0.289).

By contrast, BDNF^+/−^mice exhibited choice behavior that was equivalent during the first renewal session Day 6 compared to the last EL trial on Day 5 (Figure 3B) (D6S1 vs D5S2, Holm-Sidak, *p* = 0.051). An EL effect was developed by the second session on Day 6, however (Figure 3B) (D6S2 vs D5S2, Holm-Sidak, *p* = 0.024).

Taken together, these data suggest that BDNF is required for information updating related to context-dependent EL and renewal.

### In the absence of a context change choice adaptation is slower in BDNF+/-mice compared to wt

We examined choice adaptation during the two days of EL in the AAA paradigm (D5S1-D6S2, Figure 4A). Here, significantly slower EL effects were evident in BDNF^+/−^mice compared to WT (ANOVA, F_(1,56)_ = 6.92, *p* = 0.011)

**Figure 4:**
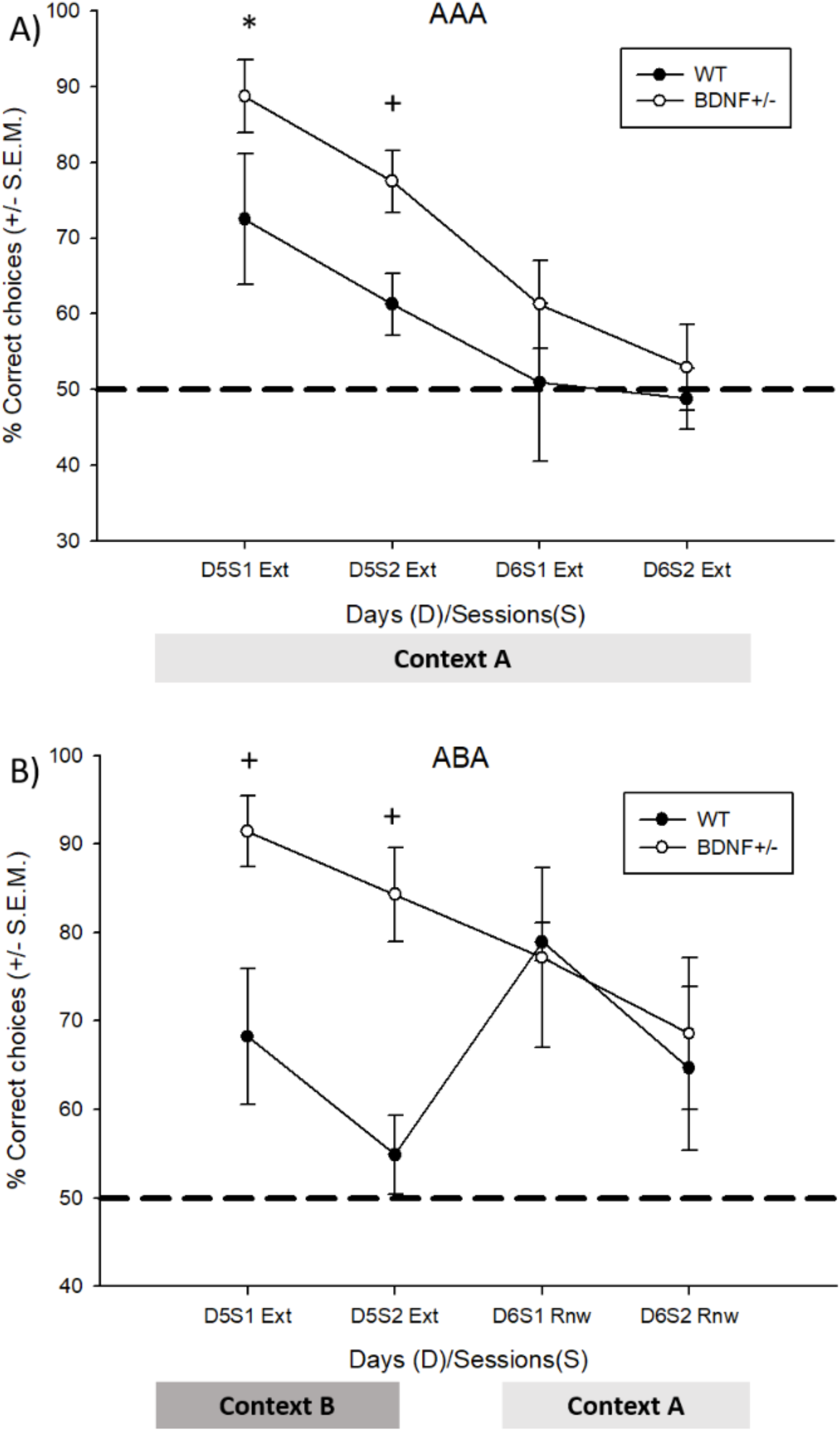
Fewer correct decisions are made by BDNF^**+/-**^ **mice during EL in both the AAA and ABA paradigms. BDNF**^**+/-**^ **mice exhibit perseverance during renewal sessions**. The graphs show the percentage of correct decisions made by the WT and BDNF^+/−^ mice during the EL sessions on Days 5 and 6 in the AAA cohort, and during EL (D5) and renewal (D6) in the ABA cohort. The black dashed line signifies chance levels. **A**. Choice adaptation behavior in BDNF^+/−^ mice is consistently poorer compared to wildtype (WT) littermates during EL in the AAA paradigm. **B**. BDNF^+/−^ mice show significantly impaired choice adaptation during EL in the ABA paradigm compared to WT littermates. During the renewal test, BDNF^+/−^ mice exhibit an absence of renewal, and perseverance in searching for a reward that is no longer available. *(p≤0.05) ^+^(p≤0.01)

### The change of context during EL accelerates choice adaptation in WT animals but does not affect the response rate in BDNF+/-mice

When EL occurred during a context change (Figure 4B), BDNF^+/−^mice showed a significantly impaired response compared to WT mice. Both groups acquired EL, as demonstrated in the effect of the session (Two way RM ANOVA, F_(3,56)_ = 2.83, *p* = 0.046), but a significant effect of the ‘group’ condition occurred (F_(1,56)_ = 9.649, *p* = 0.003). Here choices in WT mice reached approached chance levels by the second EL session on Day 5 mice (D5S2, Figure 4B). When renewal was tested on Day 6, WT mice showed a renewal effect during the first session (D6S1 vs EL, ANOVA F_(3,21)_ = 3.42, *p* = 0.031, D6S1 vs D5S2, Holm-Sidak *p* = 0.004).

These effects were different in BDNF^+/−^mice (Figure 4B). While there were differences across Day 4 to Day 6 in the BDNF^+/−^ animals, as revealed by one way rm ANOVA (F_(3,21)_ = 4.36 *p* = 0.01), a post-hoc Holm-Sidak test failed to reveal differences between D4S2 and D5S2 (*p* = 0.52), corresponding to EL, or when comparing performance during D5S2 with D6S1 (*p* = 0.61), corresponding to renewal. A significant difference in performance was found when the end of the acquisition and the end of extinction were compared (D4S2 vs D6S2) (*p* < 0.01).

## 4. DISCUSSION

This study demonstrates that hippocampal BDNF depletion is associated with an impairment of spatial appetitive EL. Whereas EL is delayed in the absence of a context-dependent change, both EL and renewal are impaired in a context-dependent EL paradigm. This study provides novel insights into the role of BDNF in appetitive EL of spatial experience, suggesting in particular that BDNF is required for information updating related to EL.

The contribution of BDNF to associative learning, and to hippocampus–dependent forms of memory is experience-dependent (Aarse et al., 2016). In line with this, spatial reference memory in the water maze task is impaired when BDNF is unavailable (Minichiello et al., 1999). Most strikingly, BDNF is required for forms of hippocampal synaptic plasticity that are *directly triggered* by spatial learning (Aarse et al., 2016). This suggests that BDNF plays a pivotal role in hippocampus-dependent experience encoding. In line with this, both BDNF^+/−^ mice and BDNF val66Met models exhibit deficiencies in contextual fear memory (Chen et al., 2006). Furthermore, in mice expressing promotor-IV of the BDNF gene, hippocampal BDNF expression is impaired and EL of context-related fear association and perseverance are changed (Sakata et al., 2013; Sakata and Duke, 2014). Our study extends these insights into the role of BDNF in EL, by adding novel findings with regard to the involvement of BDNF in appetitive EL of a spatial task.

Spatial appetitive EL recruits hippocampal gene encoding (Mendez-Couz et al., 2019). As yet, it is unclear to what extent this process involves hippocampal synaptic plasticity, but context-dependent forms of synaptic plasticity that are triggered by learning have been reported (Manahan-Vaughan and Braunewell, 1999; Kemp and Manahan-Vaughan, 2004). Long-term hippocampus-dependent memory is associated with the expression of persistent forms of LTP (Kemp and Manahan-Vaughan, 2004; Novkovic et al., 2015) and LTD (Kemp and Manahan-Vaughan, 2004; 2007; 2012; Kemp et al., 2013). Both learning-facilitated LTD and spatial reference memory are impaired in BDNF^+/**−**^ mice (Novkovic et al., 2015; Aarse et al., 2016), suggesting that it is critically required for the physiological encoding of hippocampus-dependent memory.

In the present study, when an ABA paradigm was used, BDNF^+/−^ mice exhibited deficits in EL and renewal. This process is likely to recruit structures other than the hippocampus: Eddy et al. (2016) proposed that there is a region of medial prefrontal cortex encompassing both dorsomedial prefrontal cortex and ventromedial prefrontal cortex that is important for ABA renewal of extinguished instrumental responding for a food reward. This study suggests that the dorsomedial prefrontal cortex plays a role in ABA renewal of extinguished operant reinforced response (lever pressing) and that inactivation of the vmPFC affected both extinction and ABA renewal expression. The vmPFC might be critical for the expression of both an excitatory and an inhibitory context-response association. This study differs from ours however, in that the focus was on appetitive operant conditioning, whereas our focus was on appetitive spatial EL.

The hippocampus is involved in appetitive spatial EL in an ABA paradigm (Mendez-Couz and Manahan-Vaughan, 2019). Peters et al. (2010) have proposed that BDNF may be transported from the hippocampus to the infralimbic cortex to facilitate the extinction of fear, as the hippocampus projects to the infralimbic cortex (Hoover and Vertes, 2007). Rosas-Vidal et al. (2018) reported a BDNF expression increase in the ventral hippocampus, but not in the infralimbic cortex, following fear conditioned extinction training. This may be explained by the likelihood that inactivation of the prelimbic and infralimbic areas of the mPFC impairs memory for multiple task switches, but not for the flexible selection of familiar tasks (Rich and Shapiro, 2007). Furthermore, flexible spatial learning depends on both the dorsal and ventral hippocampus and their functional interactions with the mPFC (Avigan et al., 2020).

Although we do not yet know to what extent the prefrontal cortex participates in spatial appetitive EL, the results of the present study indicate a differentiated role for BDNF depending on whether EL is supported by a change in context, or not. Here, we detected no overall change in EL if it occurred in the absence of a context change, although the establishment of EL was delayed. This is surprising given that EL in a spatial appetitive AAA paradigm requires the activation of beta-adrenergic receptors, indicating that it involves more attentive effort. It also requires the involvement of mGlu5 receptors, suggesting that it may involve dendritic protein synthesis (Huber et al., 2005). One possibility, that is as yet, unexplored, is that EL of spatial appetitive learning in the absence of a context change does not require the involvement of the hippocampus. In line with this possibility, when AAA EL was carried out with regard to a hidden platform task in a water maze, the prelimbic and infralimbic areas of mPFC showed a higher c-Fos activation, although the dorsal hippocampus did not (Mendez-Couz et al, 2014). Recent studies show that the recruitment of a specific neural system into EL is also determined by the protocol and kind of memory undergoing EL. Here, the hippocampus and dorsolateral striatum are presumed to mediate different kinds of EL, similar to their roles in the acquisition or consolidation of implicit and explicit tasks (see: Goodman and Packard 2019). In our study, despite the delay in achieving the same level of EL as in wt mice, mice effectively extinguished the original learning response. The fact that this can occur despite BDNF depletion suggests that synaptic remodeling plays a subordinate role in this process.

By contrast, BDNF was intrinsically involved in spatial appetitive EL that was supported by a context-change. This is a process that requires activation of dopamine D1/D5 receptors (André and Manahan-Vaughan, 2016) and the triggering of immediate early gene expression (Mendez-Couz and Manahan-Vaughan, 2019) supporting the likelihood that *de novo* protein synthesis is involved. Furthermore, in our experiment, the BDNF^+/−^ mice exhibited an absence of renewal behavior, although their acquisition was unimpaired. One possibility is that this effect is derived from the requirement of BDNF in the memory retrieval process. This suggests that the EL experience during the change of context in BDNF^+/−^ mice undermined the stability of the previously encoded memory of the T-maze task. This not only suggests that context-dependent EL involves information updating, rather than *de novo* encoding, but it also indicates that both processes require BDNF-mediated cell signaling. Previously, we reported no change in proBDNF levels in BDNF^+/−^ mice (Novkovic et al., 2015). Thus, the likely candidate in this process is the TrkB-mediated receptor pathway that is activated preferentially by mature BDNF (Lee et al., 2001; Ibánez, 2002; Teng et al., 2005) and leads to activation of MAP kinase and phosphatidylinositol-3-kinase (PI3K) (Kaplan and Miller, 2000; Patapoutian and Reichardt, 2001; Sweat, 2001; Huang and Reichardt, 2003). This signaling pathway has been proposed to enable the integration of newborn cells into hippocampal networks (Bergami et al., 2008) that then improve the robustness of a stored representation. The BDNF/TRkB pathway is also intrinsically involved in hippocampal synaptic plasticity and memory (Snyder et al., 2001; Bruel-Jungerman et al., 2007; Sisti et al., 2007).

### Conclusions

We report here, that EL during a context-dependent change is more sensitive to the reduced availability of BDNF than EL in the absence of a context change. In the latter, AAA paradigm, EL was delayed, whereas, in the former ABA paradigm, EL was potently impaired. Both impairments occurred despite the fact that acquisition learning was unaffected by BDNF knockdown. This means that novel spatial learning *per se* can occur in BDNF^+/−^ mice, but information updating related to EL is disrupted. In the ABA paradigm, renewal is also impaired in BDNF^+/−^ mice. In fact, animals persisted in looking for the reward in the target arm, even though it was no longer present (and had been removed prior to the EL trials), indicating that inappropriate perseverance occurred. Taken together these findings suggest that BDNF is required for learning flexibility and information updating.

## Data Availability Statement

The data that support the findings of this study are available from the corresponding author upon reasonable request.

## Ethics Statement

The animal study was reviewed and approved by Landesamt für Arbeitsschutz, Naturschutz, Umweltschutz und Verbraucherschutz, Nordrhein Westfalen, Germany.

## Author contributions

The study was designed by DM-V. Experiments were conducted by MM-C and analyzed by both authors. MM-C and DM-V wrote the paper.

## Acknowledgments

Authors thank Petra Küsener for technical assistance, Sebastian Wenzlaff and Anne-Katrin Reker for behavioral assistance, and Nadine Kollosch for animal care.

## References

Aarse, J., Herlitze, S., and Manahan-Vaughan, D. (2016). The requirement of BDNF for hippocampal synaptic plasticity is experience-dependent. Hippocampus 26(6), 739–751. doi: 10.1002/hipo.22555.

Andre, M.A., Gunturkun, O., and Manahan-Vaughan, D. (2014). The metabotropic glutamate receptor, mGlu5, is required for extinction learning that occurs in the absence of a context change. Hippocampus 25(2), 149–158. doi: 10.1002/hipo.22359.

André, M.A., Gunturkun, O., and Manahan-Vaughan, D. (2015a). The metabotropic glutamate receptor, mGlu5, is required for extinction learning that occurs in the absence of a context change. Hippocampus 25(2), 149–158. doi: 10.1002/hipo.22359.

André, M.A., and Manahan-Vaughan, D. (2015). Involvement of Dopamine D1/D5 and D2 Receptors in Context-Dependent Extinction Learning and Memory Reinstatement. Front Behav Neurosci 9, 372. doi: 10.3389/fnbeh.2015.00372.

André, M.A., Wolf, O.T., and Manahan-Vaughan, D. (2015b). Beta-adrenergic receptors support attention to extinction learning that occurs in the absence, but not the presence, of a context change. Front Behav Neurosci 9, 125. doi: 10.3389/fnbeh.2015.00125.

Avigan, P.D., Cammack, K., and Shapiro, M.L. (2020). Flexible spatial learning requires both the dorsal and ventral hippocampus and their functional interactions with the prefrontal cortex. Hippocampus 30(7), 733–744. doi: 10.1002/hipo.23198.

Bath, K.G., Schilit, A., and Lee, F.S. (2013). Stress effects on BDNF expression: effects of age, sex, and form of stress. Neuroscience 239, 149–156. doi: 10.1016/j.neuroscience.2013.01.074.

Bouton, M.E. (2019). Extinction of instrumental (operant) learning: interference, varieties of context, and mechanisms of contextual control. Psychopharmacology (Berl) 236(1), 7–19. doi: 10.1007/s00213-018-5076-4.

Bouton, M.E., and Bolles, R.C. (1979). Role of conditioned contextual stimuli in reinstatement of extinguished fear. Journal of Experimental Psychology: Animal Behavior Processes 5(4), 368.

Brunoni, A.R., Lopes, M., and Fregni, F. (2008). A systematic review and meta-analysis of clinical studies on major depression and BDNF levels: implications for the role of neuroplasticity in depression. Int J Neuropsychopharmacol 11(8), 1169–1180. doi: 10.1017/S1461145708009309.

Chen, Z.Y., Jing, D., Bath, K.G., Ieraci, A., Khan, T., Siao, C.J., et al. (2006). Genetic variant BDNF (Val66Met) polymorphism alters anxiety-related behavior. Science 314(5796), 140–143. doi: 10.1126/science.1129663.

Eddy, M.C., Todd, T.P., Bouton, M.E., and Green, J.T. (2016). Medial prefrontal cortex involvement in the expression of extinction and ABA renewal of instrumental behavior for a food reinforcer. Neurobiol Learn Mem 128, 33–39. doi: 10.1016/j.nlm.2015.12.003.

Goh J, Manahan-Vaughan D (2013) Spatial Object Recognition Enables Endogenous LTD that Curtails LTP in the Mouse Hippocampus. Cerebral Cortex 23:1118–1125. doi: 10.1093/cercor/bhs089.

Goodman, E. and Packard (2019) There Is More Than One Kind of Extinction Learning.Front. Sys Neurosci. https://doi.org/10.3389/fnsys.2019 https://doi.org/10.3389/fnsys.2019.00016.0001

Hoang TH, Aliane V, Manahan-Vaughan D (2018) Novel exploration of positional or directional spatial cues induces Arc mRNA expression in different hippocampal subfields: evidence for parallel information processing and the “what” stream. Hippocampus, 28: 315-326.doi: 10.1002/hipo.22833

Hofer, M.M., and Barde, Y.A. (1988). Brain-derived neurotrophic factor prevents neuronal death in vivo. Nature 331(6153), 261–262. doi: 10.1038/331261a0.

Hoover, W.B., and Vertes, R.P. (2007). Anatomical analysis of afferent projections to the medial prefrontal cortex in the rat. Brain Struct Funct 212(2), 149–179. doi: 10.1007/s00429-007-0150-4.

Hopkins, M.E., and Bucci, D.J. (2010). BDNF expression in perirhinal cortex is associated with exercise-induced improvement in object recognition memory. Neurobiol Learn Mem 94(2), 278–284. doi: 10.1016/j.nlm.2010.06.006.

Karpova, N.N., Pickenhagen, A., Lindholm, J., Tiraboschi, E., Kulesskaya, N., Agustsdottir, A., et al. (2011). Fear erasure in mice requires synergy between antidepressant drugs and extinction training. Science 334(6063), 1731–1734. doi: 10.1126/science.1214592.

Kemp, A., and Manahan-Vaughan, D. (2004). Hippocampal long-term depression and long-term potentiation encode different aspects of novelty acquisition. Proc Natl Acad Sci U S A 101(21), 8192–8197. doi: 10.1073/pnas.0402650101.

Kemp, A., and Manahan-Vaughan, D. (2007). Hippocampal long-term depression: master or minion in declarative memory processes? Trends Neurosci 30(3), 111–118. doi: 10.1016/j.tins.2007.01.002.

Kemp, A., and Manahan-Vaughan, D. (2012). Passive spatial perception facilitates the expression of persistent hippocampal long-term depression. Cereb Cortex 22(7), 1614–1621. doi: 10.1093/cercor/bhr233.

Kemp, A., Tischmeyer, W., and Manahan-Vaughan, D. (2013). Learning-facilitated long-term depression requires activation of the immediate early gene, c-fos, and is transcription dependent. Behav Brain Res 254, 83–91. doi: 10.1016/j.bbr.2013.04.036.

Kirtley, A., and Thomas, K.L. (2010). The exclusive induction of extinction is gated by BDNF. Learn Mem 17(12), 612–619. doi: 10.1101/lm.1877010.

Korte, M., Carroll, P., Wolf, E., Brem, G., Thoenen, H., and Bonhoeffer, T. (1995). Hippocampal long-term potentiation is impaired in mice lacking brain-derived neurotrophic factor. Proc Natl Acad Sci U S A 92(19), 8856–8860. doi: 10.1073/pnas.92.19.8856.

Linnarsson, S., Bjorklund, A., and Ernfors, P. (1997). Learning deficit in BDNF mutant mice. Eur J Neurosci 9(12), 2581–2587. doi: 10.1111/j.1460-9568.1997.tb01687.x.

Lu, H., Park, H., and Poo, M.M. (2014). Spike-timing-dependent BDNF secretion and synaptic plasticity. Philos Trans R Soc Lond B Biol Sci 369(1633), 20130132. doi: 10.1098/rstb.2013.0132.

Lucidi-Phillipi, C.A., Gage, F.H., Shults, C.W., Jones, K.R., Reichardt, L.F., and Kang, U.J. (1995). Brain-derived neurotrophic factor-transduced fibroblasts: production of BDNF and effects of grafting to the adult rat brain. J Comp Neurol 354(3), 361–376. doi: 10.1002/cne.903540306.

Ma, J., Zhang, Z., Su, Y., Kang, L., Geng, D., Wang, Y., et al. (2013). Magnetic stimulation modulates structural synaptic plasticity and regulates BDNF-TrkB signal pathway in cultured hippocampal neurons. Neurochem Int 62(1), 84–91. doi: 10.1016/j.neuint.2012.11.010.

Ma, Y.L., Wang, H.L., Wu, H.C., Wei, C.L., and Lee, E.H. (1998). Brain-derived neurotrophic factor antisense oligonucleotide impairs memory retention and inhibits long-term potentiation in rats. Neuroscience 82(4), 957–967. doi: 10.1016/s0306-4522(97)00325-4.

Manahan-Vaughan, D., and Braunewell, K.H. (1999). Novelty acquisition is associated with induction of hippocampal long-term depression. Proc Natl Acad Sci U S A 96(15), 8739–8744. doi: 10.1073/pnas.96.15.8739.

Mendez-Couz, M., Becker, J.M., and Manahan-Vaughan, D. (2019). Functional Compartmentalization of the Contribution of Hippocampal Subfields to Context-Dependent Extinction Learning. Front Behav Neurosci 13, 256. doi: 10.3389/fnbeh.2019.00256.

Mendez-Couz, M., Conejo, N.M., Vallejo, G., and Arias, J.L. (2015). Brain functional network changes following Prelimbic area inactivation in a spatial memory extinction task. Behav Brain Res. doi: 10.1016/j.bbr.2015.03.033.

Mendez-Couz, M., Gonzalez-Pardo, H., Vallejo, G., Arias, J.L., and Conejo, N.M. (2016). Spatial memory extinction differentially affects dorsal and ventral hippocampal metabolic activity and associated functional brain networks. Hippocampus 26(10), 1265–1275. doi: 10.1002/hipo.22602.

Minichiello, L., Korte, M., Wolfer, D., Kuhn, R., Unsicker, K., Cestari, V., et al. (1999). Essential role for TrkB receptors in hippocampus-mediated learning. Neuron 24(2), 401–414. doi: 10.1016/s0896-6273(00)80853-3.

Notaras, M., and van den Buuse, M. (2019). Brain-Derived Neurotrophic Factor (BDNF): Novel Insights into Regulation and Genetic Variation. Neuroscientist 25(5), 434–454. doi: 10.1177/1073858418810142.

Notaras, M., and van den Buuse, M. (2020). Neurobiology of BDNF in fear memory, sensitivity to stress, and stress-related disorders. Mol Psychiatry. doi: 10.1038/s41380-019-0639-2.

Novkovic, T., Mittmann, T., and Manahan-Vaughan, D. (2015). BDNF contributes to the facilitation of hippocampal synaptic plasticity and learning enabled by environmental enrichment. Hippocampus 25(1), 1–15. doi: 10.1002/hipo.22342.

Park, H., and Poo, M.M. (2013). Neurotrophin regulation of neural circuit development and function. Nat Rev Neurosci 14(1), 7–23. doi: 10.1038/nrn3379.

Peters, J., Dieppa-Perea, L.M., Melendez, L.M., and Quirk, G.J. (2010). Induction of fear extinction with hippocampal-infralimbic BDNF. Science 328(5983), 1288–1290. doi: 10.1126/science.1186909.

Poo, M.M. (2001). Neurotrophins as synaptic modulators. Nat Rev Neurosci 2(1), 24–32. doi: 10.1038/35049004.

Psotta, L., Lessmann, V., and Endres, T. (2013). Impaired fear extinction learning in adult heterozygous BDNF knock-out mice. Neurobiol Learn Mem 103, 34–38. doi: 10.1016/j.nlm.2013.03.003.

Rich, E.L., and Shapiro, M.L. (2007). Prelimbic/infralimbic inactivation impairs memory for multiple task switches, but not flexible selection of familiar tasks. J Neurosci. 27(17), 4747–4755. doi: 10.1523/jneurosci.0369-07.2007.

Rosas-Vidal, L.E., Lozada-Miranda, V., Cantres-Rosario, Y., Vega-Medina, A., Melendez, L., and Quirk, G.J. (2018). Alteration of BDNF in the medial prefrontal cortex and the ventral hippocampus impairs extinction of avoidance. Neuropsychopharmacology 43(13), 2636–2644. doi: 10.1038/s41386-018-0176-8.

Sakata, K., and Duke, S.M. (2014). Lack of BDNF expression through promoter IV disturbs expression of monoamine genes in the frontal cortex and hippocampus. Neuroscience 260, 265–275. doi: 10.1016/j.neuroscience.2013.12.013.

Sakata, K., Martinowich, K., Woo, N.H., Schloesser, R.J., Jimenez, D.V., Ji, Y., et al. (2013). Role of activity-dependent BDNF expression in hippocampal-prefrontal cortical regulation of behavioral perseverance. Proc Natl Acad Sci U S A 110(37), 15103–15108. doi: 10.1073/pnas.1222872110.

Soliman, F., Glatt, C.E., Bath, K.G., Levita, L., Jones, R.M., Pattwell, S.S., et al. (2010). A genetic variant BDNF polymorphism alters extinction learning in both mouse and human. Science 327(5967), 863–866. doi: 10.1126/science.1181886.

Todd, T.P., Vurbic, D., and Bouton, M.E. (2014). Mechanisms of renewal after the extinction of discriminated operant behavior. J Exp Psychol Anim Learn Cogn 40(3), 355–368. doi: 10.1037/xan0000021.

Wiescholleck, V., André, M.A., and Manahan-Vaughan, D. (2014). Early age-dependent impairments of context-dependent extinction learning, object recognition, and object-place learning occur in rats. Hippocampus 24(3), 270–279. doi: 10.1002/hipo.22220.

Zagrebelsky, M., and Korte, M. (2014). Form follows function: BDNF and its involvement in sculpting the function and structure of synapses. Neuropharmacology 76 Pt C, 628–638. doi: 10.1016/j.neuropharm.2013.05.029.

